# SARS-CoV-2 originated from SARS-CoV-1-related Bat-CoVs through Pan-CoVs rather than from SARS-CoV-2-related Bat-CoVs

**DOI:** 10.1101/2021.09.06.459210

**Authors:** Perumal Arumugam Desingu, K. Nagarajan

## Abstract

The emergence of the novel SARS-CoV-2 in 2019 sparked a dispute concerning its origin. Here, we report that the SARS-CoV-2 originated through pangolin-coronavirus (Pan-CoVs) from the SARS-CoV-related-bat-coronaviruses (SARS-CoV-1-rB-CoVs) rather than from SARS-CoV-2-related-bat-coronaviruses (SARS-CoV-2-rB-CoVs), in contrast to the previous thought. Further, our analyses strongly suggest that the Pan-CoVs evolved from the SARS-CoV-1-rB-CoVs without recombination. Further, our results suggest that the SARS-CoV-1-rB-CoVs’ perhaps jumped into the pangolin, which forced the viruses to mutate and adapt to the new host, and resulted in the origin of Pan-CoVs. Surprisingly, the Pan-CoVs formed an evolutionary intermediate between SARS-CoV-2 and SARS-CoV-2-rB-CoVs at the spike gene. Our findings also suggest that the Pan-CoV/GX and Pan-CoV/Guangdong lineages recombined to form the SARS-CoV-2 spike gene. We also found evidence that the SARS-CoV-2-rB-CoVs spike gene evolved via recombination between Pan-CoV/Guangdong and SARS-CoV-1-rB-CoVs. Overall, our findings suggest that the SARS-CoV-2 emerged from SARS-CoV-1-rB-CoVs through host jumping.

## Introduction

SARS-CoV-2, a readily human adaptable, transmissible, and infective virus, first surfaced in China in December 2019 and quickly spread over the world (1-3), posing significant evolutionary questions about its genesis. The majority of the investigations, on the other hand, focused on the SARS-like CoV-2’s genesis from SARS-CoV-2-rB-CoVs (4-7) through Pan-CoVs (7-9) via genetic recombination (8,10-13). Furthermore, SARS-CoV-2 is also the seventh coronavirus to infect humans, and it is closely linked to SARS-CoVs (SARS-CoV-1). Although SARS-CoV-2 and related bat coronaviruses (SARS-CoV-2-rB-CoVs) have been identified (2,4-7), none of them have developed an evolutionary intermediate connecting link between SARS-CoV-1/SARS-CoV-1-rB-CoVs and SARS-CoV-2/SARS-CoV-2-rB-CoVs across their genome (8,10-13). Similarly, when compared to SARS-CoV-2, all the SARS-CoV-2-rB-CoVs, except RaTG13, displayed a lot of genetic heterogeneity in the spike gene (5-7,14), which is critical for defining host range, transmissibility, virus entry, human-human infection, and host-mediated immune responses (15-19). In light of this, the evolution of SARS-CoV-2-rB-CoVs in bats from SARS-CoV-1-rB-CoVs, as well as the origin of SARS-CoV-2 from SARS-CoV-1 is unknown.

The SARS-CoV-2 is designated as a new/novel coronavirus due to its genetic diversity in the pp1a, spike protein, ORF3, and protein 8 genes (1,3,4). The CoVs in general, the spike protein influences the host range, virus transmission, attachment, and entry (15-19); pp1a (nsp3, largest gene of pp1a) which determines the viral replication/transcription complex (RTC) and also regulates various other functions such as ssRNA binding, nucleocapsid binding, de-MARylation, dePARylation, ADPr binding, G-quadruples binding, protease, and deISGylation (20-27); ORF3 is a gene that controls virus release via viral ion channels (viroporins) (28,29), and regulate autophagy to favour viral replication (30-32); and ORF8 (protein 8) controls antigen presentation through MHC-I and regulates host immune surveillance (33,34). Moreover, in the pp1a, ORF3, and protein 8 areas, the SARS-CoV-2-rB-CoVs shared a close genetic link with the SARS-CoV-2. When compared to SARS-CoV-2, the SARS-CoV-2-rB-CoVs (except RaTG13) revealed a lot of genetic variation in the spike gene. Overall, the origins of the novel pp1a, ORF3, protein 8, and spike genes in SARS-CoV-2/SARS-CoV-2-rB-CoVs are unknown.

In this study, we found evidence that the SARS-CoV-2 evolved from SARS-CoV-1/SARS-CoV-1-rB-CoVs through Pan-CoVs as an evolutionary intermediate between SARS-CoV-1/SARS-CoV-1-rB-CoVs and SARS-CoV-2/SARS-CoV-2-rB-CoVs across their genome without recombination events. Our data further suggest that SARS-CoV-2 arose through the recombination of two different Pan-CoVs lineages. We also present evidence that the SARS-CoV-2-rB-CoVs are the result of recombination between Pan-CoVs and SARS-CoV-1-rB-CoVs.

## Results

### Pan-CoVs genetic diversity at the complete genome levels

In our phylogenetic, net between-group mean distance (NBGMD), and similarity plot-based analysis, the SARS-CoV-2-rB-CoVs of RaTG13, RShSTT200, RShSTT182, RacCS203, RmYN02, and RpYN06 formed a close genetic link with the SARS-CoV-2 (**Figure 1A, 1B and 1C**). On the other hand, in our phylogenetic tree analysis, we also observed that the SARS-CoV-2-rB-CoVs of bat/PrC31, bat-SL-CoVZC45, and bat-SL-CoVZXC21 viruses constituted out-group from SARS-CoV-2 (**Figure 1A**). Consistent with these results, our NBGMD analysis revealed that these viruses (bat/PrC31, bat-SL-CoVZC45, and bat-SL-CoVZXC21) had displayed around 10% genetic diversity when compared to SARS-CoV-2 (**Figure 1B**). Furthermore, in our similarity plot-based analysis, the bat/PrC31, bat-SL-CoVZC45, and bat-SL-CoVZXC21 revealed a great deal of variability in the pp1b gene when compared to SARS-CoV-2 (**Figure 1C**).

**Figure 1.**
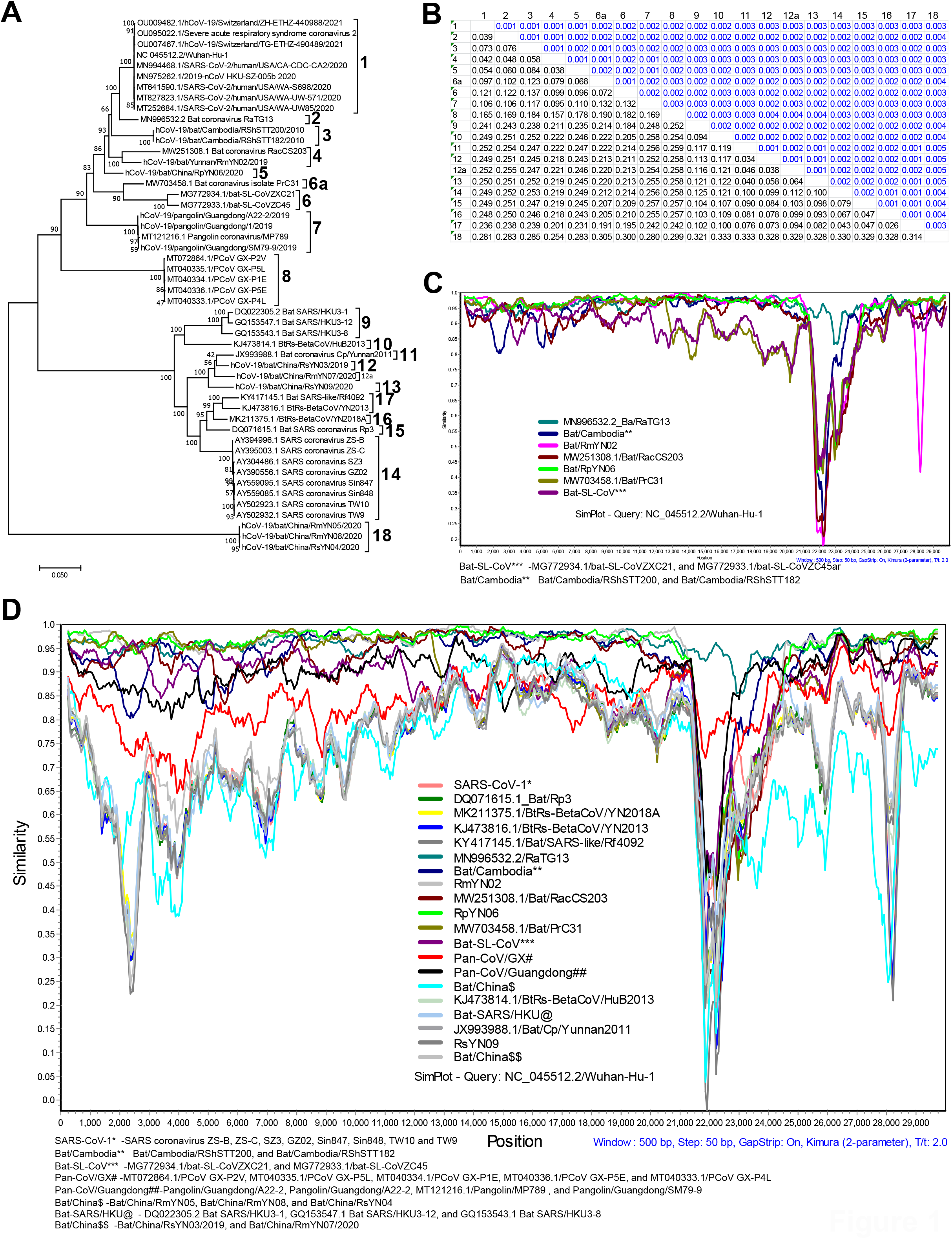
SARS-CoV-2 and Pan-CoVs genetic diversity at the complete genome. (**A &B**) The Phylogenetic tree (A) and net between-group mean distance (B) respectively for the Pan-CoVs, SARS-CoV-2, and related viruses using the complete genome sequence. (**C**) Similarity plot analysis using the complete genomes of the SARS-CoV-2 and SARS-CoV-2-rB-CoVs. SARS-CoV-2 is set as a query sequence (**D**) Similarity plot analysis using the complete genomes of the SARS-CoV-2, SARS-CoV-2-rB-CoVs, Pan-CoVs, SARS-CoV-1, and SARS-CoV-1-rB-CoVs. SARS-CoV-2 is set as a query sequence.

Next, in our phylogenetic analysis, the Pan-CoVs are divided into two lineages: Pan-CoV/Guangdong and Pan-CoV/GX (**Figure 1A**), and it is also supported by NBGMD (**Figure 1B**) and similarity plot-based analysis (**Figure 1D; Supplementary Figure 1A-1B**). Surprisingly, these Pan-CoVs lineages formed an evolutionary bridge between SARS-CoV-1/SARS-CoV-1-rB-CoVs and SARS-CoV-2/SARS-CoV-2-rB-CoVs throughout their genome (**Figure 1D; Supplementary Figure 1A-1B**). The Pan-CoV/Guangdong lineage developed a closer relationship with the SARS-CoV-2 and SARS-CoV-2-rB-CoVs than the Pan-CoV/GX lineage in the phylogenetic tree (**Figure 1A**) and also in NBGMD analyses (**Figure 1B**). In our similarity plot-based analysis, the Pan-CoV/Guangdong lineage shared a stronger genetic resemblance with the SARS-CoV-2/SARS-CoV-2-rB-CoVs except for the spike gene S1-N-terminal domain (S1-NTD) (**Figure 1D; Supplementary Figure 1A-1B**). Remarkably in our further analysis, the Pan-CoV/GX showed an evolutionary intermediate between the SARS-CoV-1/SARS-CoV-1-rB-CoVs and SARS-CoV-2/SARS-CoV-2-rB-CoVs throughout their genome (**Figure 1D; Supplementary Figure 1A-1B**). Collectively, these findings suggest that the SARS-CoV-1/SARS-CoV-1-rB-CoVs jumped into pangolins and evolved into Pan-CoV/GX in pangolins as a result of host selection and adaption pressure. The Pan-CoV/GX in the pangolin further evolved into the Pan-CoV/Guangdong lineage, which is strikingly similar to the SARS-CoV-2/SARS-CoV-2-rB-CoVs due to probable better host adaptation and immune evasion selection pressure.

### Evolution of pp1a in SARS-CoV-2 from SARS-CoV-1 through Pan-CoVs

According to earlier studies, the pp1a gene was reported as a novel gene in SARS-CoV-2 (1,3,4). To explore the evolutionary genesis of the pp1a gene, we used the SARS-CoV-2 nucleotide sequences from the first nucleotide to 11725nt (nucleotide numbering according to NC 045512.2/Wuhan-Hu-1). Because the SARS-CoV-1 viruses displayed a lot of diversity from 1-11725nt and displayed a closer identity from 11726-21561nt with the SARS-CoV-2/SARS-CoV-2-rB-CoVs, therefore we divided the pp1ab into two different regions for the analysis as follows: 1-11725nt and 11726-21561nt for analysis (**Figure 1D; Supplementary Figure 1A-1B**). The SARS-CoV-2-rB-CoVs of bat-SL-CoVZC45 and bat-SL-CoVZXC21 have nearly identical sequences to the SARS-CoV-1 from the 11725nt to the end of the pp1ab gene (**Figure 1D; Supplementary Figure 1A-1B**). The SARS-CoV-2-rB-CoVs of the RpYN06, RmYN02, and Bat/PrC31 viruses formed a closer relationship with SARS-CoV-2 in this region (1-11725nt) than the RaTG13 virus (**Figure 2A-2C; Supplementary Figure 2A-2B**). Surprisingly in this location, the Pan-CoVs formed an evolutionary bridge between SARS-CoV-1/SARS-CoV-1-rB-CoVs and SARS-CoV-2/SARS-CoV-2-rB-CoVs (**Figure2A-2C; Supplementary Figure 2A-2B)**.

**Figure 2.**
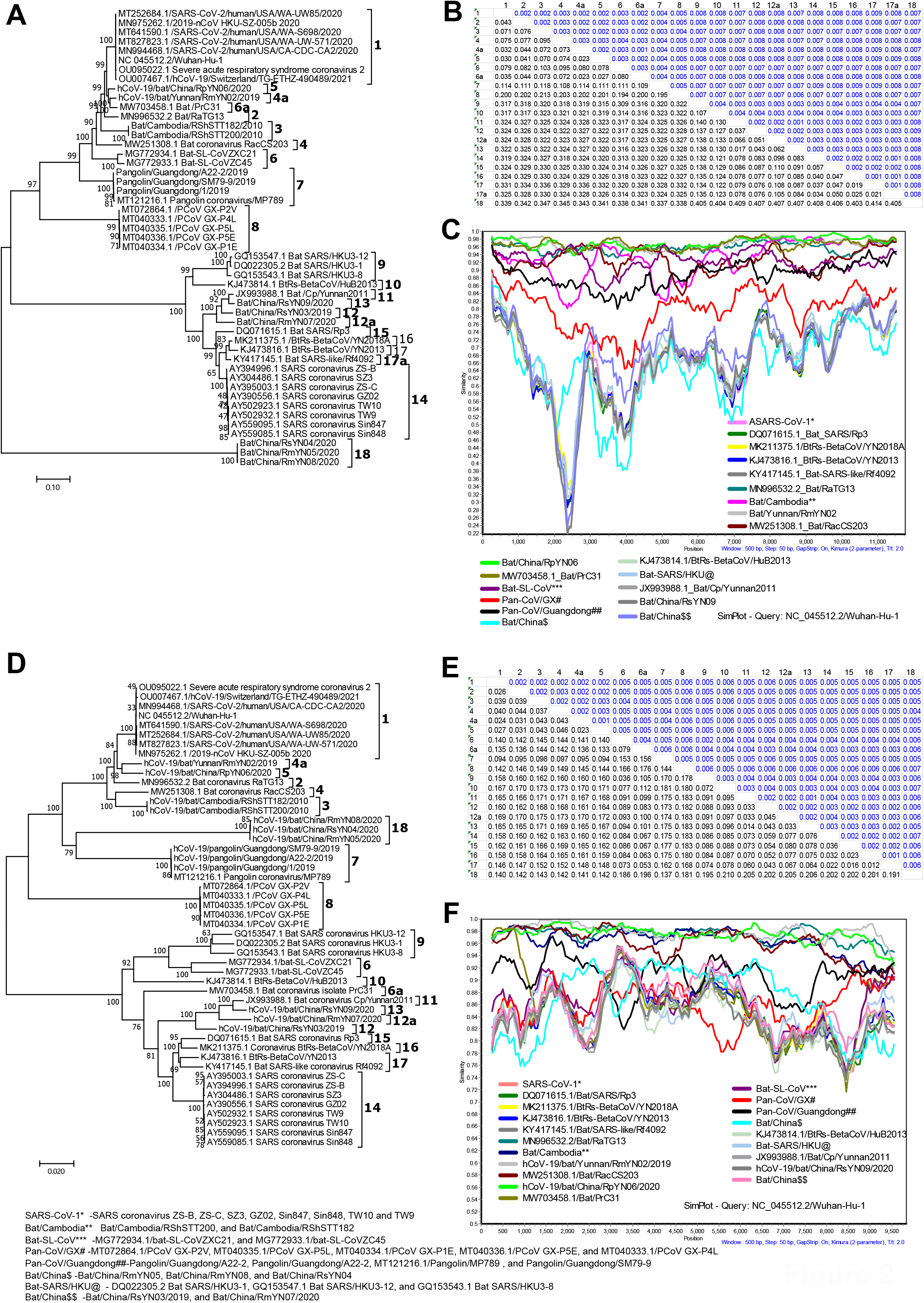
The genetic diversity analysis of the pp1ab gene. (**A-C**) The Phylogenetic tree (A), net between-group mean distance (B), and similarity plot analysis (C) respectively for the Pan-CoVs, SARS-CoV-2, and related viruses using part of pp1ab (1-11725nt). SARS-CoV-2 is set as a query sequence. (**D-F**) The Phylogenetic tree (D), net between-group mean distance (E), and similarity plot analysis (F) respectively for the Pan-CoVs, SARS-CoV-2, and related viruses using part of pp1ab from 11726nt to 21561 nucleotides. SARS-CoV-2 is set as a query sequence.

Next, in comparison to the previous region, the genetic diversity between SARS-CoV-1 and SARS-CoV-2 is lowest in the pp1ab regions of 11726nt to 21561 nucleotides (**Figure 2D-2F; Supplementary Figure 3A-3B)**. Surprisingly, the bat/PrC31, bat-SL-CoVZC45, and bat-SL-CoVZXC21 SARS-CoV-2-rB-CoVs had nearly identical sequence similarity to the SARS-CoV-1-rB-CoVs of bat-SARS-coronavirus-HKU3 viruses (**Figure 2D-2F; Supplementary Figure 3A-3B)**. Remarkably, Pan-CoVs also created an evolutionary intermediate between SARS-CoV-1/SARS-CoV-1-rB-CoVs and SARS-CoV-2/SARS-CoV-2-rB-CoVs in this region (**Figure 2D-2F; Supplementary Figure 3A-3B)**. Overall, our findings imply that throughout the pp1ab gene, Pan-CoVs formed an evolutionary intermediate between SARS-CoV-1 and SARS-CoV-2. Altogether, our results suggest that the Pan-CoVs gained this pp1a gene from SARS-CoV-1/SARS-CoV-1-rB-CoVs and it might have evolved into a novel pp1a gene in pangolin through new host adaptation and immune evasion mediated evolution. Further, the jumping of Pan-CoVs into humans/other related hosts possibly evolved into the SARS-CoV-2-specific pp1a which is seen in the ongoing outbreak. On the other hand, the SARS-CoV-2-rB-CoVs (bat/PrC31, bat-SL-CoVZC45, and bat-SL-CoVZXC21) potentially gained this pp1ab gene by recombination of Pan-CoVs and the SARS-CoV-1-rB-CoVs of bat-SARS-coronavirus-HKU3 viruses.

### Origin of novel ORF3 in SARS-CoV-2 from SARS-CoV-1 through Pan-CoVs

We investigated the evolutionary history of the novel ORF3 in SARS-CoV-2 and SARS-CoV-2-rB-CoVs in light of recent studies that suggested they acquired it (1,3,4). In this location, the Pan-CoV/GX lineage formed an evolutionary link between SARS-CoV-1 and SARS-CoV-2, while the Pan-CoV/Guangdong lineage shared a high degree of sequence similarity with SARS-CoV-2 (**Figure 3A-3C; Supplementary Figure 4A-4B**). It was surprising to find that SARS-CoV-2-rB-CoVs of bat/PrC31, bat-SL-CoVZC45, and bat-SL-CoVZXC21 viruses differed considerably from SARS-CoV-2 and other SARS-CoV-2-rB-CoVs (**Figure 3A-3C; Supplementary Figure 4A-4B**). These results indicated that as similar to the pp1a gene, the novel ORF3 also evolved in the SARS-CoV-2 from the SARS-CoV-1 through the Pan-CoVs by the host jump mediated virus adaptation and evolution.

**Figure 3.**
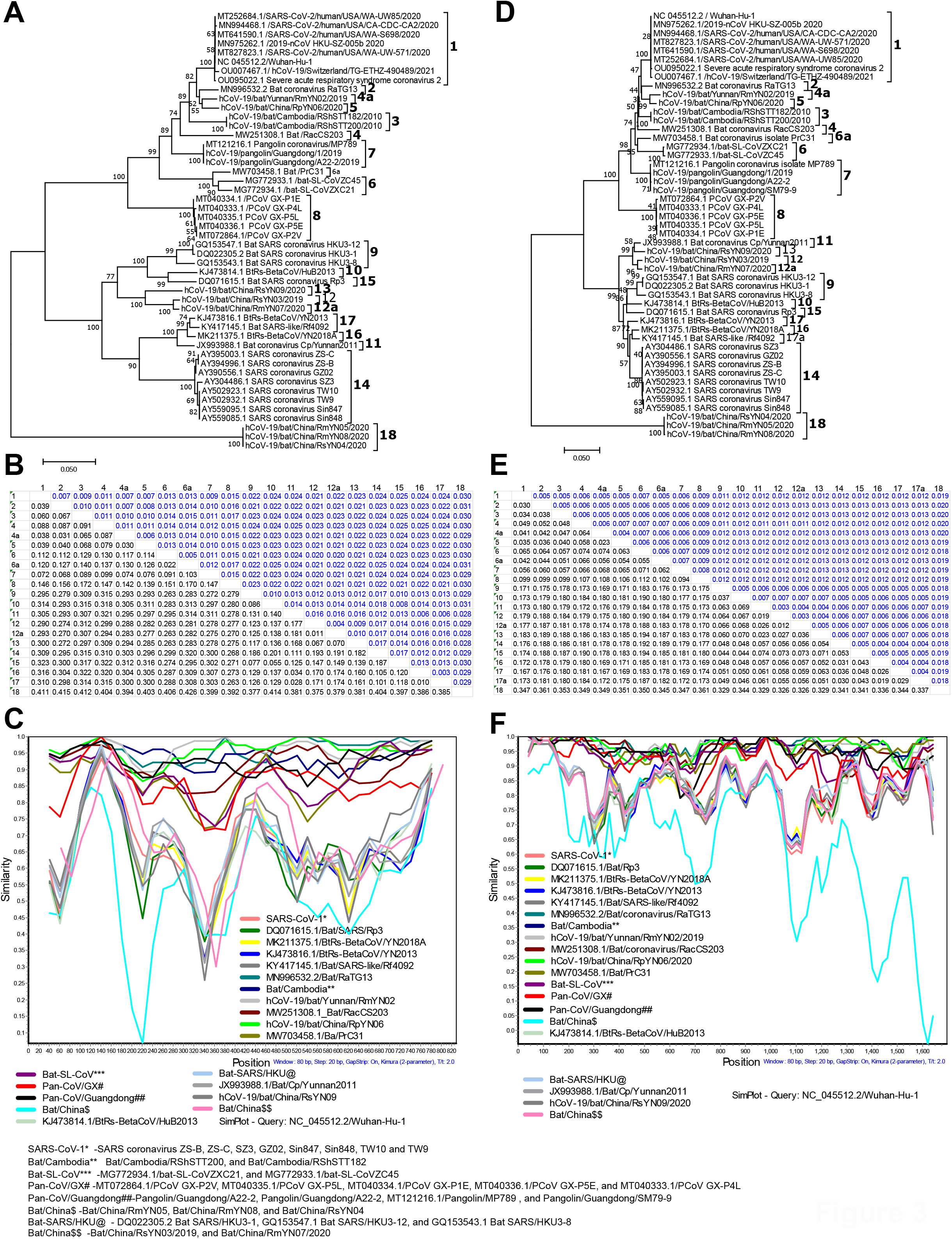
The genetic diversity analysis of ORF3, E, M, protein 6, protein 7a, and protein 7b. (**A-C**) The Phylogenetic tree (A), net between-group mean distance (B), and similarity plot analysis (C) respectively for the Pan-CoVs, SARS-CoV-2, and related viruses using nucleotide sequences of complete ORF3. (**D-F**) The Phylogenetic tree (D), net between-group mean distance (E), and similarity plot analysis (F) respectively for the Pan-CoVs, SARS-CoV-2, and related viruses using the nucleotide sequences of E, M, protein 6, protein 7a, and protein 7b. SARS-CoV-2 is set as a query sequence.

### Genetic diversity of the envelope (E), membrane (M), protein 6, protein 7a, and protein 7b

Next, the envelope (E), membrane (M), protein 6, protein 7a, and protein 7b portions of the Pan-CoV/Guangdong, SARS-CoV-2, and SARS-CoV-2-rB-CoVs had a significant degree of sequence similarity, whereas the Pan-CoV/GX lineage lay between the SARS-CoV-1 and SARS-CoV-2 (**Figure 3D-3F; Supplementary Figure 5A-5B**). These results suggest that the SARS-CoV-2 and other SARS-CoV-2-rB-CoVs gained E, M, protein 6, protein 7a, and protein 7b from the SARS-CoV-1 via the Pan-CoV lineages.

### Gain of novel Protein 8 in SARS-CoV-2 from SARS-CoV-1 through Pan-CoVs

Although it is widely known that SARS-CoV-2 and SARS-CoV-2-rB-CoVs have the unique protein 8 gene in their genomes (1,3,4), the evolutionary origin of this gene remains unknown. We observed that similar to ORF3, the SARS-CoV-2-rB-CoVs of bat-SL-CoVZC45 and bat-SL-CoVZXC21 viruses showed significant genetic diversity from SARS-CoV-2 and other SARS-CoV-2-rB-CoVs in the protein-8 region (**Figure 4A-4B**). Remarkably, SARS-CoV-2-rB-CoVs of bat/PrC31 on the other hand demonstrated a stronger genetic identity with SARS-CoV-2 and other SARS-CoV-2-rB-CoVs. More importantly, the Pan-CoV/Guangdong lineage shared a high level of sequence similarity with SARS-CoV-2, whereas the Pan-CoV/GX lineage was an evolutionary bridge between SARS-CoV-1 and SARS-CoV-2 (**Figure 4A-4B**). These results collectively suggest that similar to pp1a and ORF3, the SARS-CoV-2 gained the novel protein 8 from the SARS-CoV-1 through the Pan-CoVs by the host jump mediated virus adaptation and evolution.

**Figure 4.**
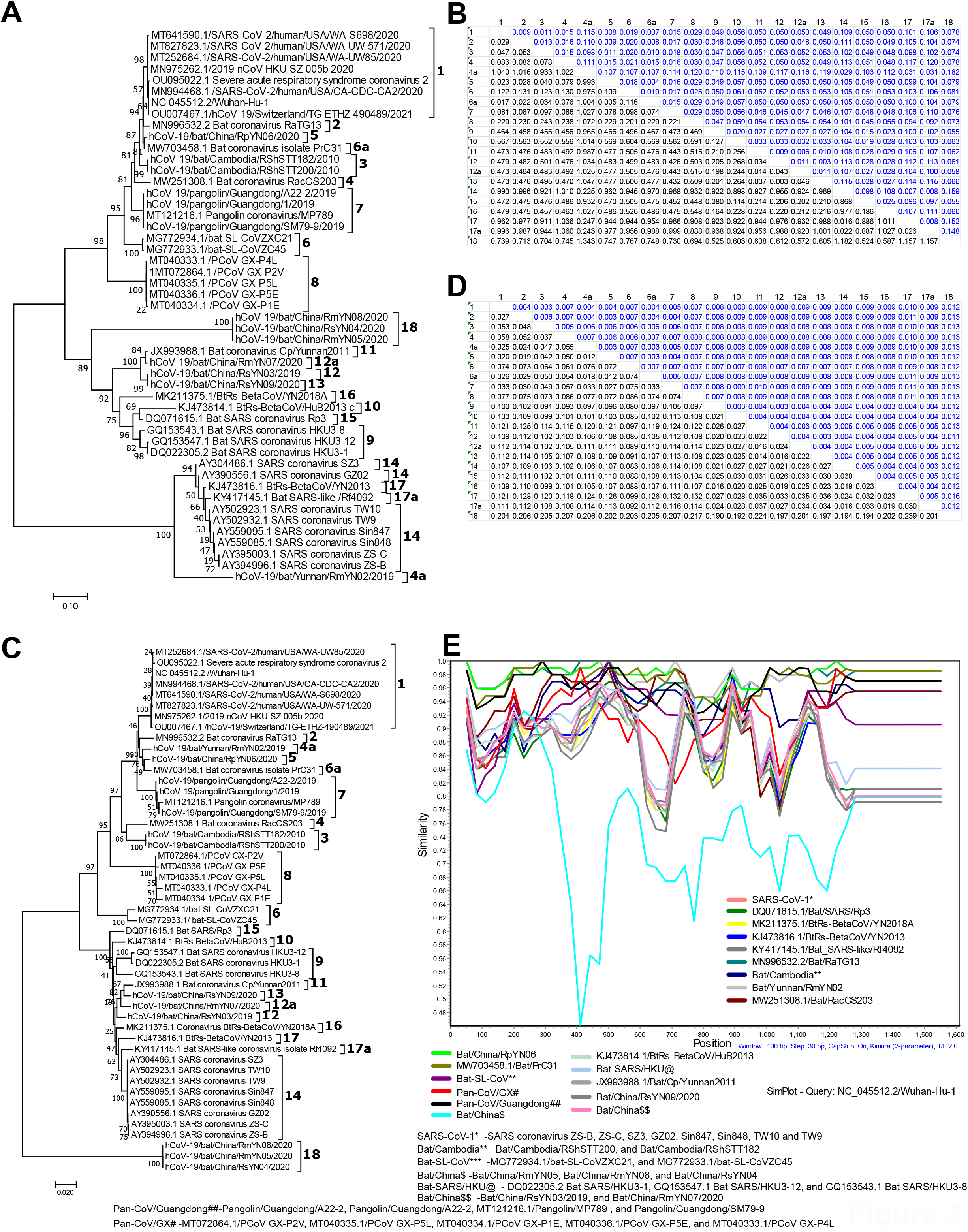
The genetic diversity analysis of ORF8 (protein 8), nucleocapsid (N), ORF10, and 3’UTR. (**A-B**) The Phylogenetic tree (A), and net between-group mean distance (B) respectively, for the Pan-CoVs, SARS-CoV-2, and related viruses using nucleotide sequences of complete ORF8 (protein 8). (**C-E**) The Phylogenetic tree (C), net between-group mean distance (D), and similarity plot analysis (E) respectively for the Pan-CoVs, SARS-CoV-2, and related viruses using the nucleotide sequences of the nucleocapsid (N), ORF10, and 3’UTR. SARS-CoV-2 is set as a query sequence.

### Genetic diversity of the nucleocapsid (N), ORF10 and 3’UTR

Next, the nucleocapsid (N), ORF10 and 3’UTR region, the Pan-CoV/Guangdong, SARS-CoV-2, and SARS-CoV-2-rB-CoVs had a closer degree of sequence similarity; however, the Pan-CoV/GX lineage formed evolutionary intermediate between SARS-CoV-1 and SARS-CoV-2 in phylogenetic, NBGMD, and similarity plot-based analysis (**Figure 4C-4E; Supplementary Figure 6A-6B**). These results indicate that the SARS-CoV-2 and other SARS-CoV-2-rB-CoVs acquired the nucleocapsid (N), ORF10, and 3’UTRregion from the SARS-CoV-1 via the Pan-CoV lineages through the host jump based evolution.

### Recombination-based origin of SARS-CoV-2 spike gene from Pan-CoVs

The spike gene which is known to determine the host range, transmission, virus entry, and host immunity (15-19) were reported to display huge genetic diversity between the SARS-CoV-2 and SARS-CoV-2-rB-CoVs (5-7,14). Consistent with this, we are also found huge genetic diversity between the SARS-CoV-2 and SARS-CoV-2-rB-CoVs (**Figure 5A-5B**) in the spike gene. Interestingly in our phylogenetic and similarity plot analysis, the Pan-CoVs displayed as an evolutionary intermediate between SARS-CoV-2 and SARS-CoV-2-rB-CoVs at the entire spike gene level (**Figure 5A&5C**). Fascinatingly, we also observed that among the Pan-CoV lineages, the Pan-CoV/Guangdong lineage had a closer evolutionary link with SARS-CoV-2 (**Figure 5C-5D; Supplementary Figure 7A-7B**). In confirmation with this, at the spike protein S1-Receptor binding domain (S1-RBD) levels, Pan-CoV/Guangdong lineage showed essentially identical sequence similarity to SARS-CoV-2 (**Figure 5C-5D; Supplementary Figure 7A-7B**). In contrast, when compared to SARS-CoV-2, the Pan-CoV/Guangdong lineage revealed a lot of genetic variation in the spike protein S1-N-Terminal domains (S1-NTD) (**Figure 5C-5D; Supplementary Figure 7A-7B**). Further, our similarity plot analysis revealed that, in comparison to SARS-CoV-2, the Pan-CoV/GX lineage showed limited variation at the S1-NTD and S1-RBD (**Figure 5C-5D; Supplementary Figure 7A-7B**). These findings point to recombination between the Pan-CoV/Guangdong and Pan-CoV/GX lineages as a probable evolutionary source for the SARS-CoV-2 virus. More interestingly, our similarity plot and recombination detection program (RDP) analyses also confirmed this recombination of Pan-CoV/Guangdong and Pan-CoV/GX lineage in the S1-NTD (**Figure 5C-5F; Supplementary Figure 7A-7B**). Collectively, our results strongly suggest that the SARS-CoV-2 spike gene was evolved through recombination of the Pan-CoV/Guangdong and Pan-CoV/GX lineages. Overall, our results strongly suggest that the SARS-CoV-2 gained all the novel genes such as pp1a, spike, ORF3, and protein 8 from the SARS-CoV-1 through the Pan-CoVs by the host jump mediated virus adaptation and evolution.

**Figure 5.**
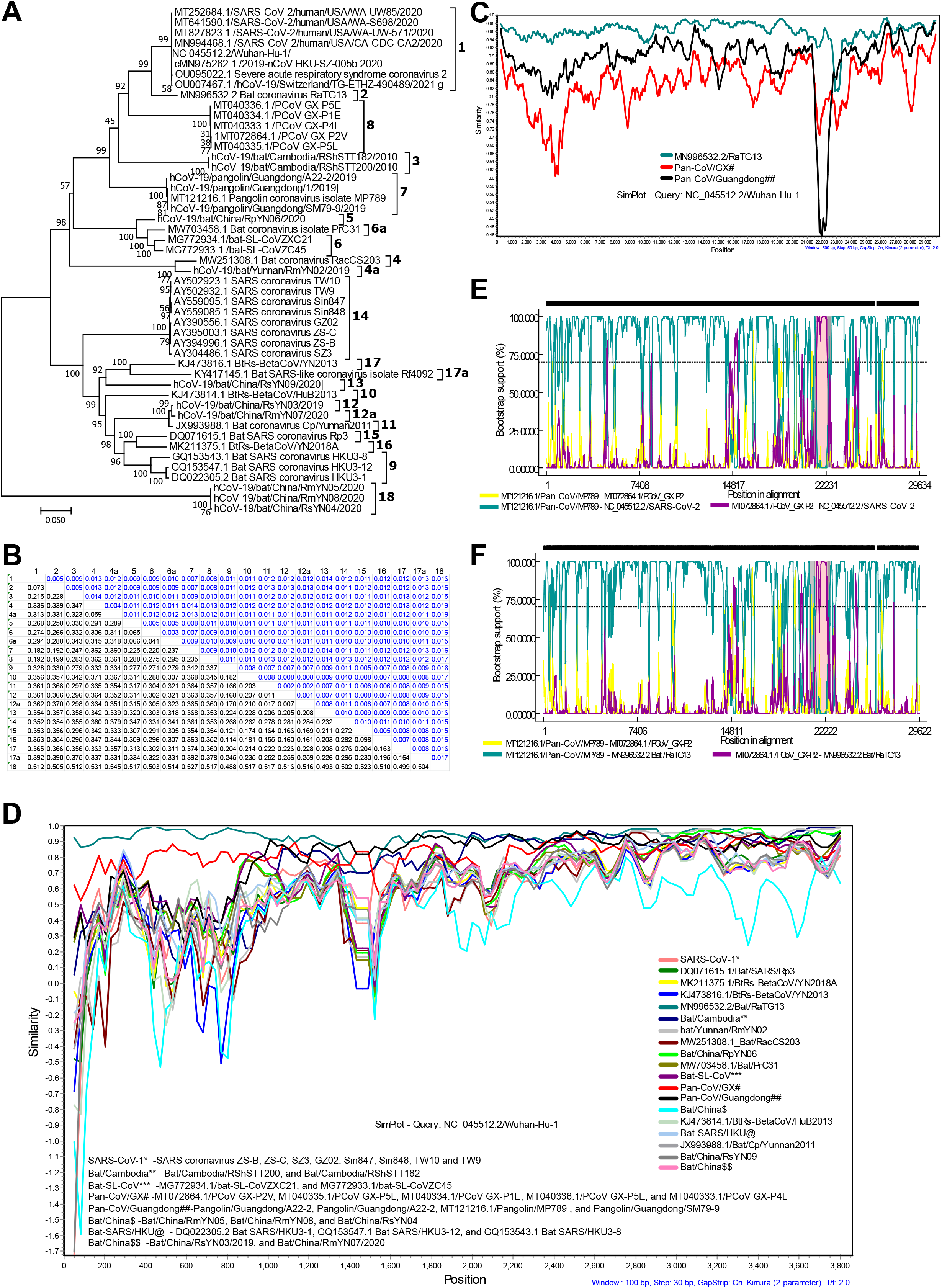
Origin SARS-CoV-2 spike gene through recombination of Pan-CoVs. (**A-B**) The Phylogenetic tree (A), and net between-group mean distance (B) respectively, for the Pan-CoVs, SARS-CoV-2, and related viruses using nucleotide sequences of complete spike gene. (**C**) Similarity plot analysis for the SARS-CoV-2, bat/RaTG13, Pan-CoV/GX, and Pan-CoV/Guangdong lineages using the complete genome nucleotide sequences. SARS-CoV-2 is used as a query sequence. (**D**) Similarity plot analysis for the Pan-CoVs, SARS-CoV-2, and related viruses using nucleotide sequences of complete spike gene. SARS-CoV-2 is used as a query sequence. (**E**) The recombination analysis with BOOTSCAN method using complete genomes of the SARS-CoV-2, PCoV-GX-P2V, and Pangolin/Guangdong/MP789. (**F**) The recombination analysis with BOOTSCAN method using complete genomes of the Bat/RaTG13, PCoV-GX-P2V, and Pangolin/Guangdong/MP789.

### Recombination-based evolution of SARS-CoV-2-rB-CoVs spike gene from Pan-CoVs

Except for RaTG13, all SARS-CoV-2-rB-CoVs had a lot of genetic variabilities when compared to SARS-CoV-2 at the entire spike (S) gene levels (5-7,14). In our phylogenetic analysis, the Pan-CoVs formed an evolutionary intermediate between the SARS-CoV-2 and SARS-CoV-2-rB-CoVs in the spike gene (**Figure 5A**). On the other hand, in NBGMD analysis, the SARS-CoV-2-rB-CoVs displayed almost equal genetic diversity with SARS-CoV-2 and SARS-CoV-1/SARS-CoV-1-rB-CoVs (**Figure 5B**). However, the SARS-CoV-2-rB-CoVs of Bat/China/RpYN06, bat/PrC31, bat-SL-CoVZC45, and bat-SL-CoVZXC21 displayed a somewhat closer identity with SARS-CoV-2 and Pan-CoVs than SARS-CoV-1/SARS-CoV-1-rB-CoVs (**Figure 5A-5B**). Further, similarity plot analysis displayed the sequence similarities between the Pan-CoV/Guangdong lineages and SARS-CoV-2-rB-CoVs (Bat/China/RpYN06, bat/PrC31, bat-SL-CoVZC45, and bat-SL-CoVZXC21) at the S1-NTD levels (**Figure 5D; Supplementary Figure 7A-7B**). Similarly, the SARS-CoV-2-rB-CoVs displayed high sequence similarity with SARS-CoV-1-rB-CoVs of bat-SARS-coronavirus-HKU3 viruses in the S1-RBD (**Figure 5D; Supplementary Figure 7A-7B**). These results indicate that the SARS-CoV-2-rB-CoVs originated through recombination of the Pan-CoV/Guangdong lineages and the SARS-CoV-1-rB-CoVs of bat-SARS-coronavirus-HKU3 viruses.

To explore the possible recombination, we performed RDP analysis. The recombination events between the Pan-CoV/Guangdong lineages and the SARS-CoV-1-rB-CoVs of bat-SARS-coronavirus-HKU3 viruses were confirmed by our analysis as follows. We characterized that the bat-SL-CoVZXC21 virus is a recombinant of Pan-CoV/Guangdong lineages with part of pp1ab and S1-RBD of SARS-CoV-1-rB-CoVs of bat-SARS-coronavirus-HKU3-1 virus (**Figure 6A-6B**). Similarly, the bat-SL-CoVZC45 virus was identified as a recombinant Pan-CoV/Guangdong lineage with the pp1ab and S1-RBD of SARS-CoV-1-rB-CoVs of bat-SARS-coronavirus-HKU3-8 virus (**Figure 6C-6D**). Furthermore, our results indicate that the SARS-CoV-2-rB-CoVs of bat/PrC31 as a recombinant Pan-CoV/Guangdong lineage with the part of pp1ab and S1-RBD of SARS-CoV-1-rB-CoVs of Bat SARS coronavirus Rp3 (**Figure 6E-6F**). Finally, our results revealed that the Bat/China/RpYN06 virus is the most closely related to SARS-CoV-2 at the complete genome level next to the bat/RaTG13 virus, which evolved through the recombination of Pan-CoV/Guangdong lineages’ and bat-SL-CoVZC45 virus (**Figure 6G-6H**). Overall, our findings strongly suggest that SARS-CoV-2-rB-CoVs arose from a series of recombination between Pan-CoVs and SARS-CoV-1-rB-CoVs.

**Figure 6.**
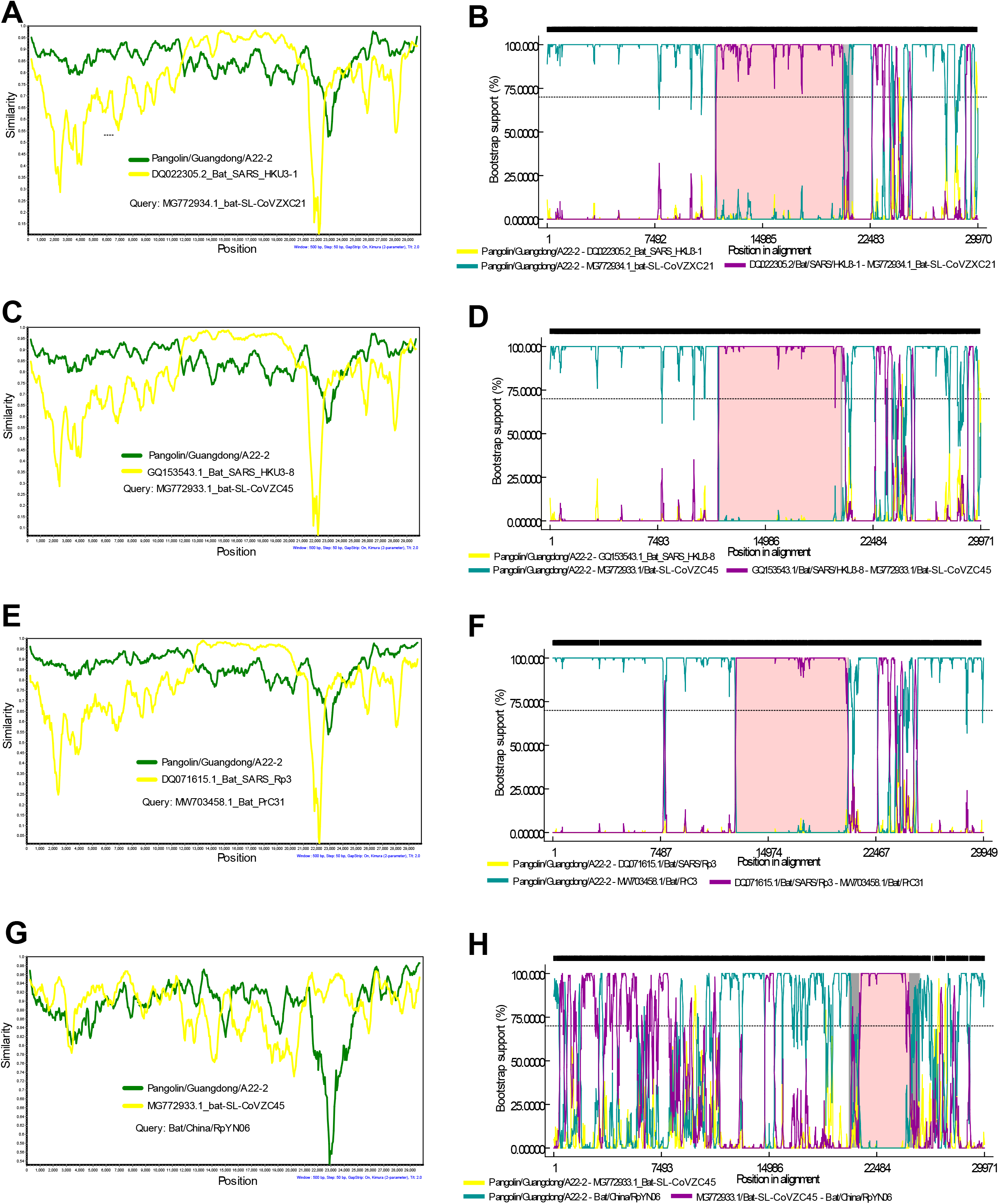
Origin SARS-CoV-2-rB-CoVs spike gene through recombination. (**A&B**) The recombination analysis with Similarity plot (A) and BOOTSCAN (B) using the complete genomes the Bat-SL-CoVZXC21, Pangolin/Guangdong/A22-2, and Bat/SARS/HKU3-1; Bat-SL-CoVZXC21 used as a query sequence. (**C&D**) The recombination analysis with Similarity plot (C) and BOOTSCAN (D) using the complete genomes the Bat-SL-CoVZC45, Pangolin/Guangdong/A22-2, and Bat/SARS/HKU3-8; Bat-SL-CoVZC45 used as a query sequence. (**E&F**) The recombination analysis: Similarity plot (E) and BOOTSCAN (F) using complete genomes of the Bat/PrC31, Pangolin/Guangdong/A22-2, and Bat/SARS/Rp3; Bat/PrC31used as a query sequence. (**G&H**) The recombination analysis with Similarity plot (G) and BOOTSCAN (H) using complete genomes of the Bat/China/RpYN06, Pangolin/Guangdong/A22-2, and Bat-SL-CoVZC45; Bat/China/RpYN06 was used as a query sequence.

## Discussion

In this study, we show that Pan-CoVs represent an intermediate stage of evolution between SARS-CoV-1/SARS-CoV-1-rB-CoVs and SARS-CoV-2/SARS-CoV-2-rB-CoVs without any recombination across their genome. Interestingly, our results indicated that the host receptor binding spike gene, the viral replication and translation controlling gene of pp1a, the virus release regulating gene of ORF3, and the host immune regulating gene of ORF8 showed considerable evolution in Pan-CoVs when compared to SARS-CoV-1/SARS-CoV-1-rB-CoVs. A virus that jumps from natural host to another must overcome the following host genetic barriers such as host surface receptors, host antiviral proteins for viral replication, virus protein translation, virus assembly, release, and the host’s innate and adaptive immune protection (35-39). Viruses used to mutate and bypass all these host barriers to accomplish in the new species through host jump (35-37,40). In majority of the time, accidental virus introduction to an unnatural host does not result in effective infection (35-37,40,41). However, in rare cases/continuous/forced exposure to an unnatural host, the viruses begin to adapt to the new host through virus mutations to tackle the host selection pressure and prolong their lifecycle (35-37,41). Viruses must then continue to evolve to increase their adaptability. Finally, it becomes highly adapted to new species to produce a large viral load that can be transmitted within the new hosts (35,37,38,40). Furthermore coronaviruses in general, the quasispecies evolutionary processes have been well described (39,42-49). To support this, the ongoing SARS-CoV-2 outbreak in humans displayed widespread mutation/deletions and evolution in the critical genes of ORF1a (46,50-54); spike gene (46,50,51,55,56); ORF3 (50,57-59); and ORF8 (33,50,60-65) those viral proteins regulates the virus transmission, better adaptability to host and evasion of the host immunity. Furthermore, the presence of selection pressure for the evolution of the SARS-CoV-2 has been highly characterized (10,52,66,67). More importantly, among the Pan-CoV/GX and Pan-CoV/Guangdong lineages, Pan-CoV/Guangdong lineage displayed high sequence identity with SARS-CoV-2 and was isolated from the clinically ill pangolins (8,9). Therefore, collectively it can be inferred that the SARS-CoV-1/SARS-CoV-1-rB-CoVs were jumped into the pangolin (new host), and then these viruses mutated/evolved in the host receptor binding spike gene, viral replication, and translation controlling gene of pp1a, virus release regulating gene of ORF3 and host immune regulating gene of ORF8 and which then evolved as Pan-CoV/GX lineage for better host adaptation. In the second stage of enhanced pangolin host adaptability, the Pan-CoV/GX lineage was then transformed to Pan-CoV/Guangdong lineage (**Figure 7**). This notion is further supported by the fact that pangolins were unwell and deceased as a result of the Pan-CoV/Guangdong lineage (8,9).

**Figure 7.**
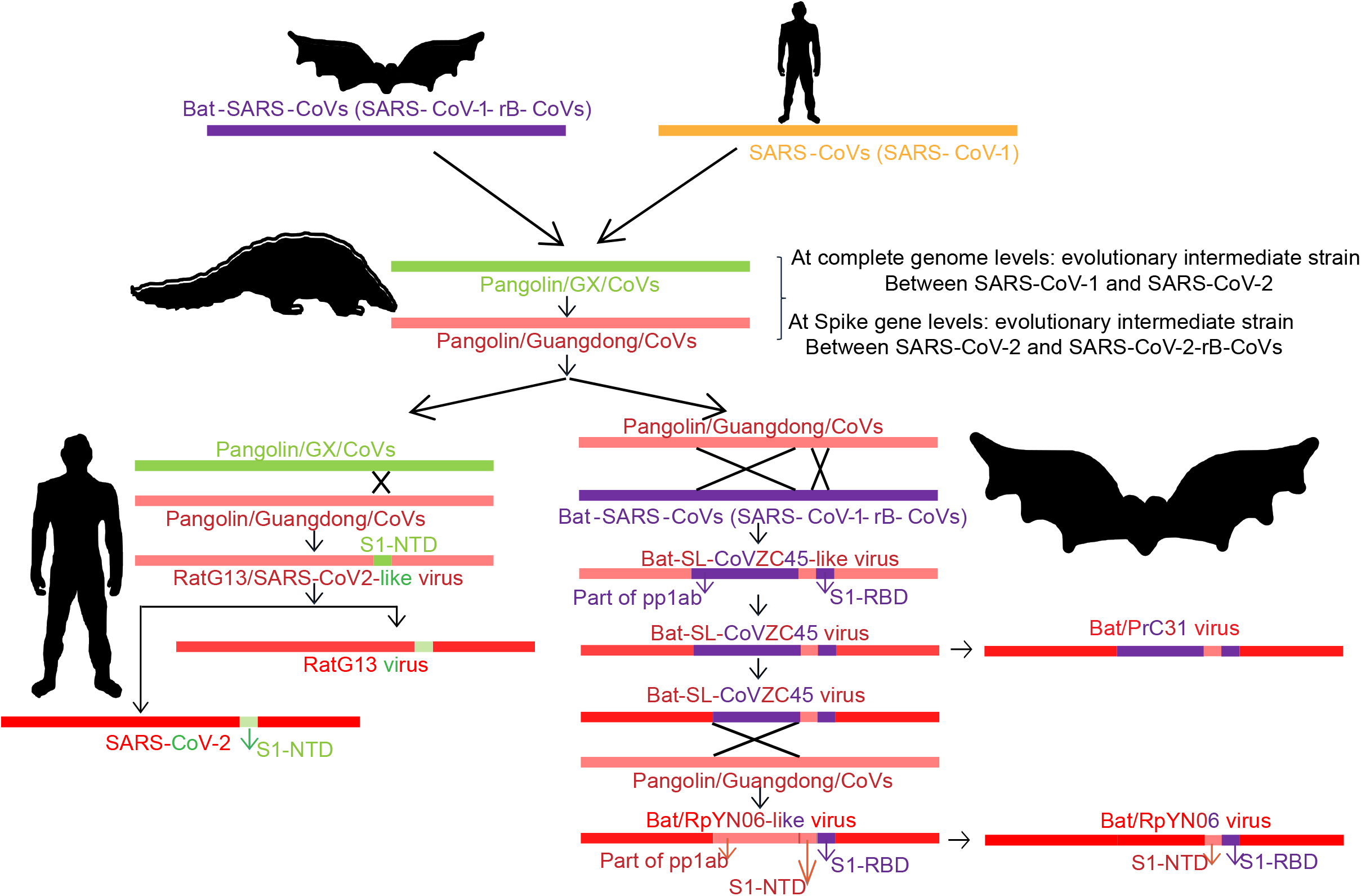
Schematic representation of origin of the SARS-CoV-2 from SARS-CoV-1 through Pan-CoVs. The SARS-CoV-1/SARS-CoV-1-rB-CoVs were perhaps jumped into the pangolin, and further evolved into a novel Pan-CoVs (two different lineages namely Pan-CoV/GX and Pan-CoV/Guangdong) by new host pressure. The SARS-CoV-2 evolved from the recombination of Pan-CoV/GX and Pan-CoV/Guangdong. Further, SARS-CoV-2-rB-CoVs were evolved through recombination of Pan-CoVs and SARS-CoV-1-rB-CoVs.

In addition, huge genetic diversity was documented between the SARS-CoV-2 and SARS-CoV-2-rB-CoVs in the spike gene (5-7,14) which is responsible for critical host range, transmissibility, and virus entry. Furthermore, the occurrence of widespread recombination events in the spike gene of SARS-CoV-2-rB-CoVs has been extensively studied for this gene’s evolution (8,10-13). As per our findings in this study, revealed that the progenitor SARS-CoV-2/RaTG13-like virus is recombinant of the Pan-CoV/Guangdong and Pan-CoV/GX lineages (**Figure 7**). In this perspective, our study implies that a recombinant of the Pan-CoV/Guangdong and Pan-CoV/GX lineages may have entered humans and begun evolving in these genes for improved adaptation (**Figure 7**). Due to the mild illness or short-chains of human-human infection, the SARS-CoV-2 viral infection may have gone unnoticed in the early stages of the virus jump (38). The virus then evolved in humans as a result of better adaptation, resulting in the SARS-CoV-2 pandemics (**Figure 7**).

Furthermore, RaTG13 is the only bat virus that shared a spike gene with SARS-CoV-2. Our findings further demonstrated that the SARS-CoV-2-rB-CoVs of bat/PrC31, bat-SL-CoVZC45, and bat-SL-CoVZXC21 were formed by recombination of Pan-CoVs and SARS-CoV-1-rB-CoVs in the part of pp1a and S1-RBD (**Figure 7**). Surprisingly, Our findings found evidence that the Bat/China/RpYN06 virus evolved by recombination of Pan-CoV/Guangdong lineages and bat-SL-CoVZC45 virus, which is the next one to bat/RaTG13 virus in terms of overall genome similarity to SARS-CoV-2 (**Figure 7**).

In conclusion, the SARS-CoV-1/SARS-CoV-1-rB-CoVs were perhaps jumped into the pangolin, further in the presence of host adaptation pressure and immune evasion, these viruses might have evolved in the critical virus entry (Spike gene), replication and translation (pp1a-nsp3), virus release (ORF3) and host immune evasion (protein 8) genes which then lead to the origin of novel Pan-CoVs. The presence of two different lineages in the Pan-CoVs indicates the different stages of host adaptation mediated evolution. The Pan-CoV/GX lineage may be originated from the first stage of host adaptation and Pan-CoV/Guangdong lineage might be the later stage of better-adapted virus in pangolins to cause disease. Further,the progenitor RaTG13/SARS-CoV-2-like virus evolved through the recombination of the Pan-CoV/GX and Pan-CoV/Guangdong lineages in the S1-NTD. Similarly, progenitor SARS-CoV-2-rB-CoVs-like viruses were evolved through recombination of Pan-CoVs and SARS-CoV-1-rB-CoVs in the part of pp1ab and S1-RBD. Further, the progenitor SARS-CoV-2-like virus and SARS-CoV-2-rB-CoVs-like virus’s gave raise to SARS-CoV-2 and SARS-CoV-2-rB-CoVs respectively through host jump mediated evolution. This study demonstrates that the SARS-CoV-2 outbreak emerged through host jump mediated virus evolution (**Figure 7**). The close monitoring and large-scale surveillance of SARS-CoV-2/SARS-CoV-2-rB-CoVs in different domestic and wild species are warranted to predict their potential to evolve as a new virus that may cause the next pandemic outbreak, and also get prepared with the vaccines and antivirals.

## Materials and Methods

### Genetic diversity analysis

#### I. Phylogenetic analysis

SARS-CoV-2 and other bat coronavirus sequences were obtained using the NCBI and GISAID databases. MEGA7 was used for the phylogenetic analysis, with the Neighbor-Joining method used to infer evolutionary history, bootstrap tests (1000 replicates), the Maximum Composite Likelihood method used to compute evolutionary distances, and gamma distribution (shape parameter = 5) used to compute the evolutionary distances being employed to model the rate variation among sites. For evolutionary comparisons, differences in composition bias among sequences were analyzed, and all ambiguous sites were discarded in the analysis for each sequence pair.

#### II. Net Between Group Mean Distance (NBGMD) analysis

The NBGMD were measured using MEGA7 tool and the Kimura 2-parameter model, with a gamma distribution (shape parameter=5) used as a model to measure rate variation among sites, a bootstrap test (1000 replicates) was used to estimate the Standard error and all ambiguous sites for each sequence pair were removed from the analysis. Above the diagonal, standard error estimates were displayed.

### Recombination analysis

#### I. SimPlot analysis

SimPlot 3.5.1 was used to calculate the percent identity between the query and reference sequences. MEGA7 was used to align the nucleotide sequences before they were exported to SimPlot 3.5.1 for further analysis. We used the Kimura two-parameter method to measure the identity between the query and reference sequences, using the 500 base pair of the window at a 50 base pair step.

#### III. Recombination Detection Program (RDP) analysis

In SARS-CoV-2 and similar viruses, RDP4 was utilized to detect possible recombination events. The nucleotides were aligned in MEGA7 before being transferred to RDP4 for further processing..For the BOOTSCAN, GENECONV, Chimaera, RDP, MaxChi, SISCAN, and 3seq methods, default parameter values were used, and a minimum of four or more approaches were examined for probable recombination using a Bonferroni adjusted p-value cut-off (0.05). The images created using the BOOTSCAN approach are shown in the figures.

## Acknowledgments

We thank Director, IISc, Bangalore, India, The Chair, Department of Microbiology and Cell Biology, Indian Institute of Science, Bengaluru, India, and Prof. Nagalingam R. Sundaresan, Department of Microbiology and Cell Biology, Indian Institute of Science, Bengaluru, India for the providing working place at IISc, Bangalore.

## Author contributions

PAD performed most of the bioinformatics experiments. KN assisted PAD for most of the bioinformatics works. PAD and KN wrote the first draft of the manuscript. PAD conceived the study, designed experiments, and wrote the final version of the manuscript.

## Data Availability

We have retrieved the nucleotide sequences from publically available NCBI and GISAID databases. Further, all the nucleotide sequences accession numbers and names are indicated in the respective figures.

## Supplementary Figures and Legends

**Supplementary Figure 1.**
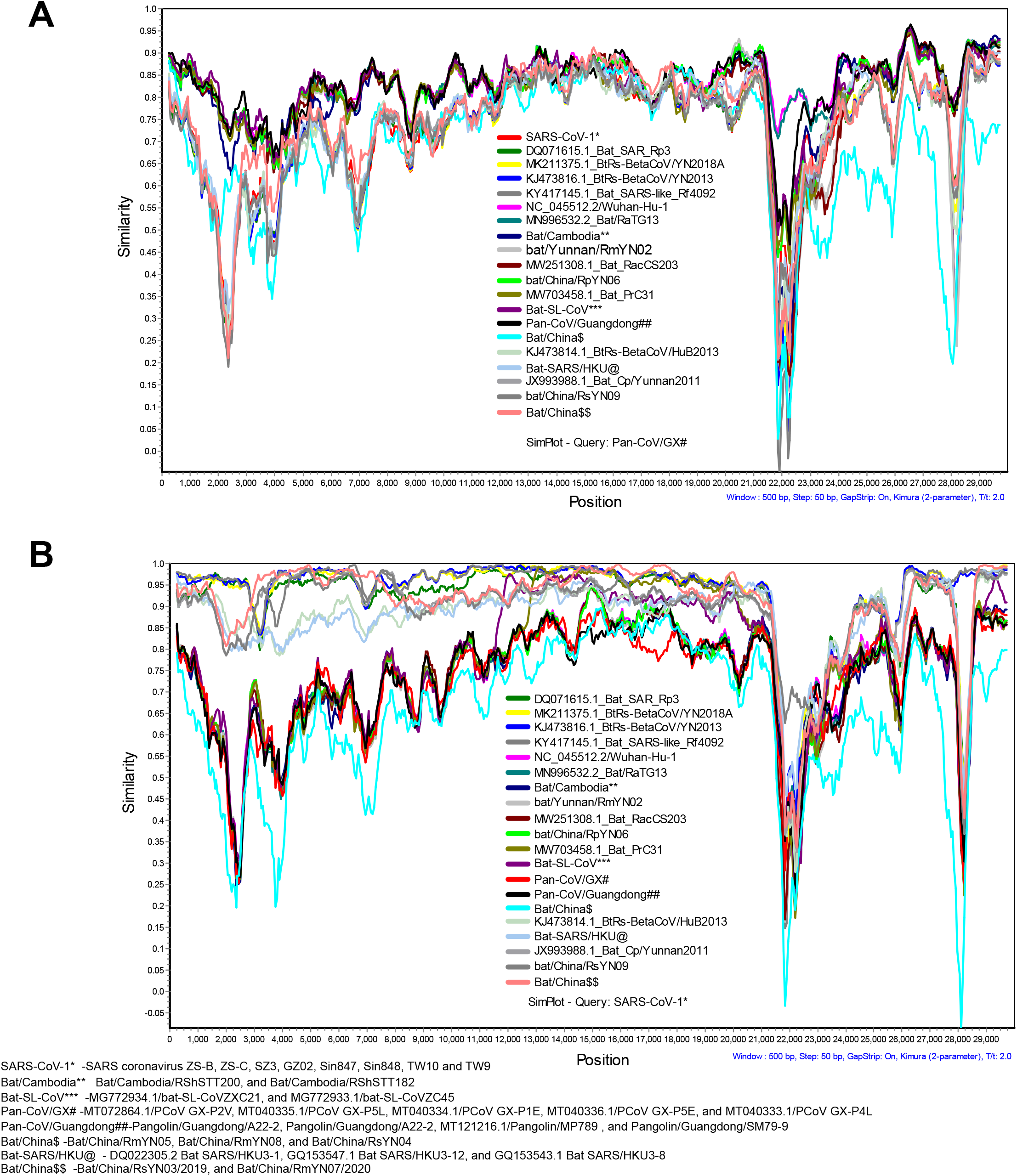
Similarity plot analysis using the complete genomes of SARS-CoV-2 and its related viruses. (**A**) Similarity plot analysis using the complete genomes of the SARS-CoV-2, SARS-CoV-2-rB-CoVs, Pan-CoVs, SARS-CoV-1, and SARS-CoV-1-rB-CoVs. The Pan-CoV/GX lineages are set as a query sequence. (**B**) Similarity plot analysis using the complete genomes of the SARS-CoV-2, SARS-CoV-2-rB-CoVs, Pan-CoVs, SARS-CoV-1, and SARS-CoV-1-rB-CoVs. The SARS-CoV-1 is set as a query sequence.

**Supplementary Figure 2.**
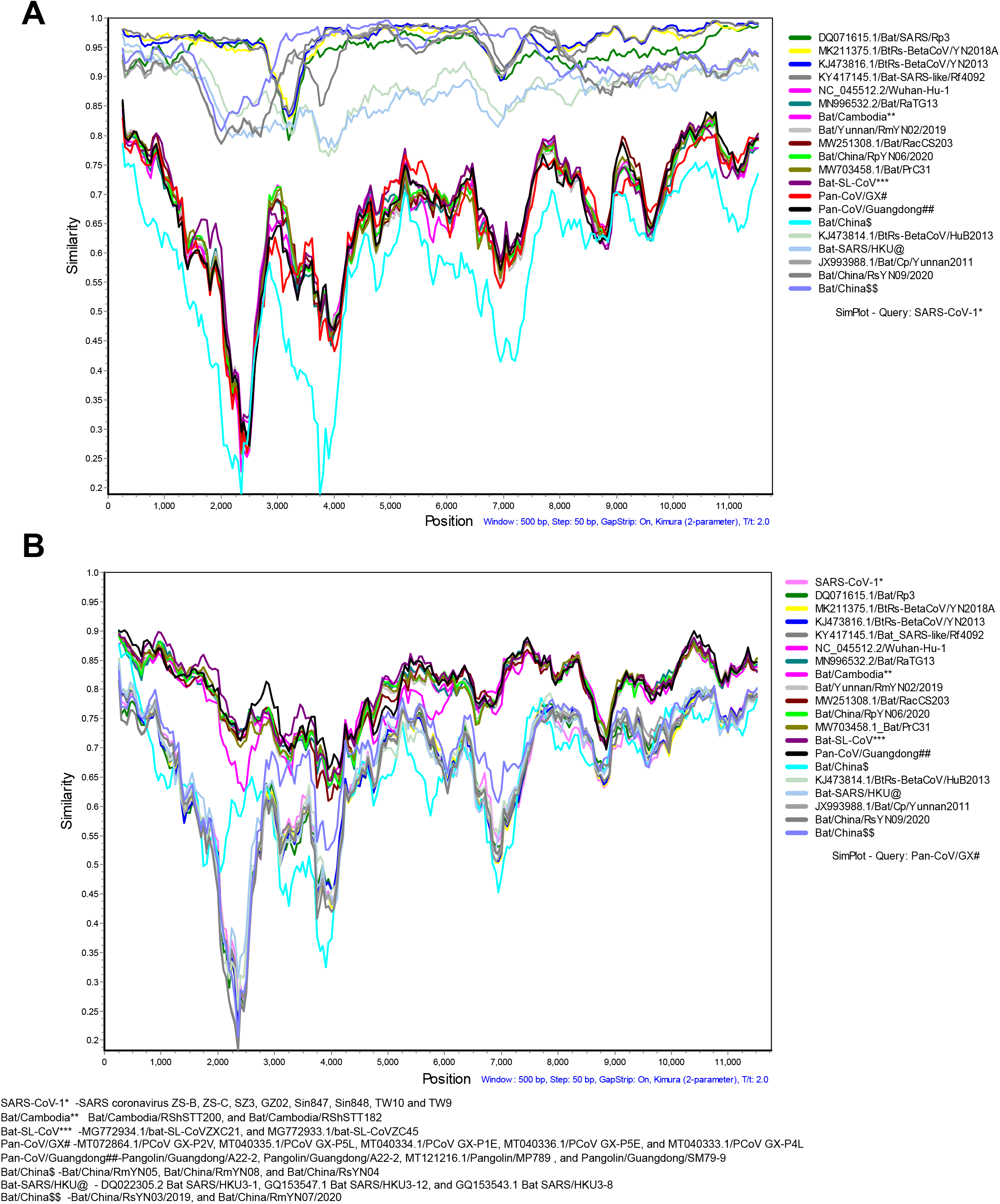
Similarity plot analysis for the part of pp1ab (1-11725nt). (**A**) Similarity plot analysis for the Pan-CoVs, SARS-CoV-2, and related viruses using part of pp1ab (1-11725nt). The SARS-CoV-1 is set as a query sequence. (**B**) Similarity plot analysis for the Pan-CoVs, SARS-CoV-2, and related viruses using part of pp1ab (1-11725nt). The Pan-CoV/GX lineages are set as a query sequence.

**Supplementary Figure 3.**
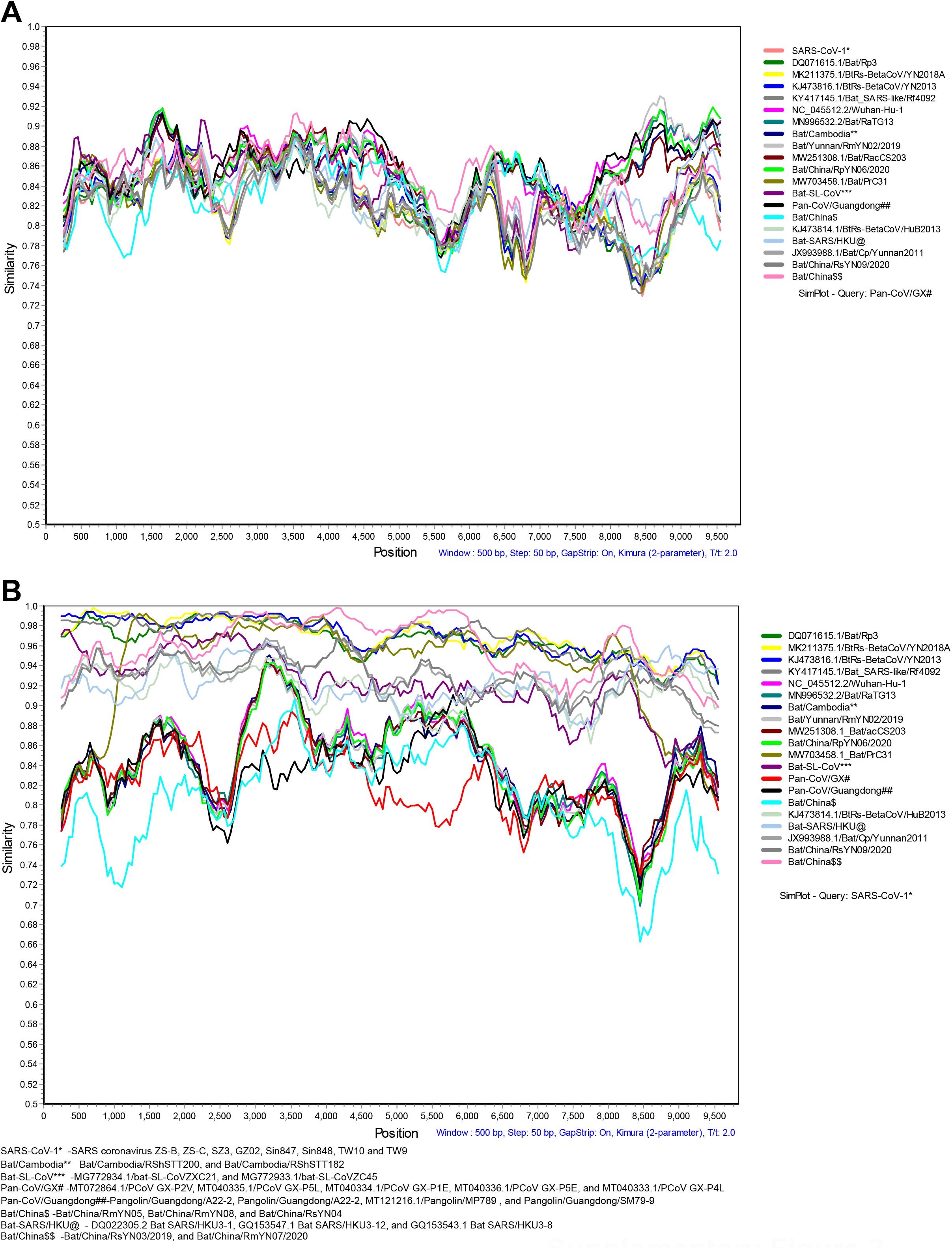
Similarity plot analysis for the part of pp1ab (11726nt to 21561nt). (**A**) Similarity plot analysis for the Pan-CoVs, SARS-CoV-2, and related viruses using part of pp1ab (11726nt to 21561nt). The Pan-CoV/GX lineages are set as a query sequence. (**B**) Similarity plot analysis for the Pan-CoVs, SARS-CoV-2, and related viruses using part of pp1ab (11726nt to 21561nt). The SARS-CoV-1 is set as a query sequence.

**Supplementary Figure 4.**
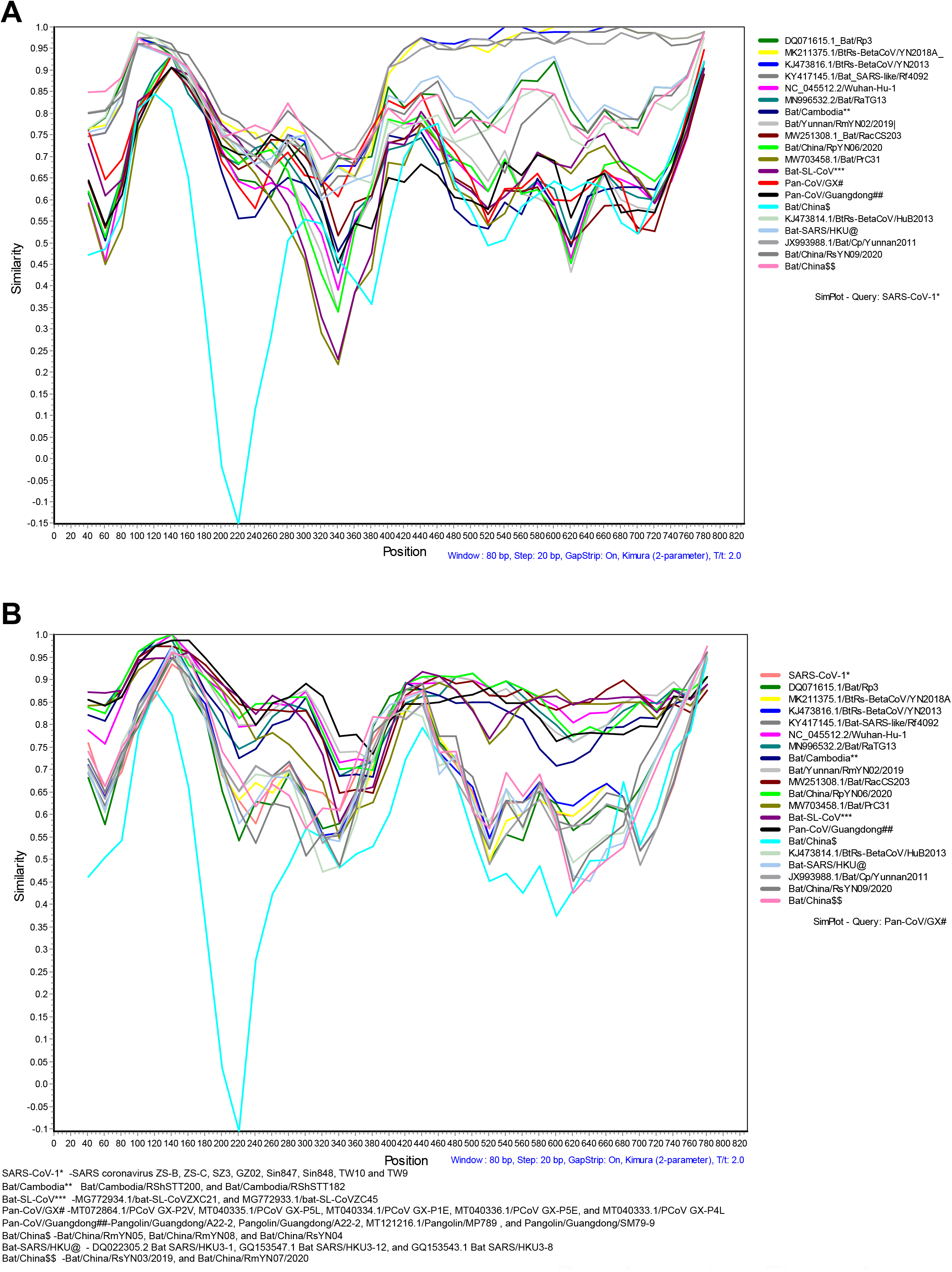
Similarity plot analysis for the ORF3. (**A**) Similarity plot analysis for the Pan-CoVs, SARS-CoV-2, and related viruses using nucleotide sequence of complete ORF3. The SARS-CoV-1 is set as a query sequence. (**B**) Similarity plot analysis for the Pan-CoVs, SARS-CoV-2, and related viruses using nucleotide sequence of complete ORF3. The Pan-CoV/GX lineages are set as a query sequence.

**Supplementary Figure 5.**
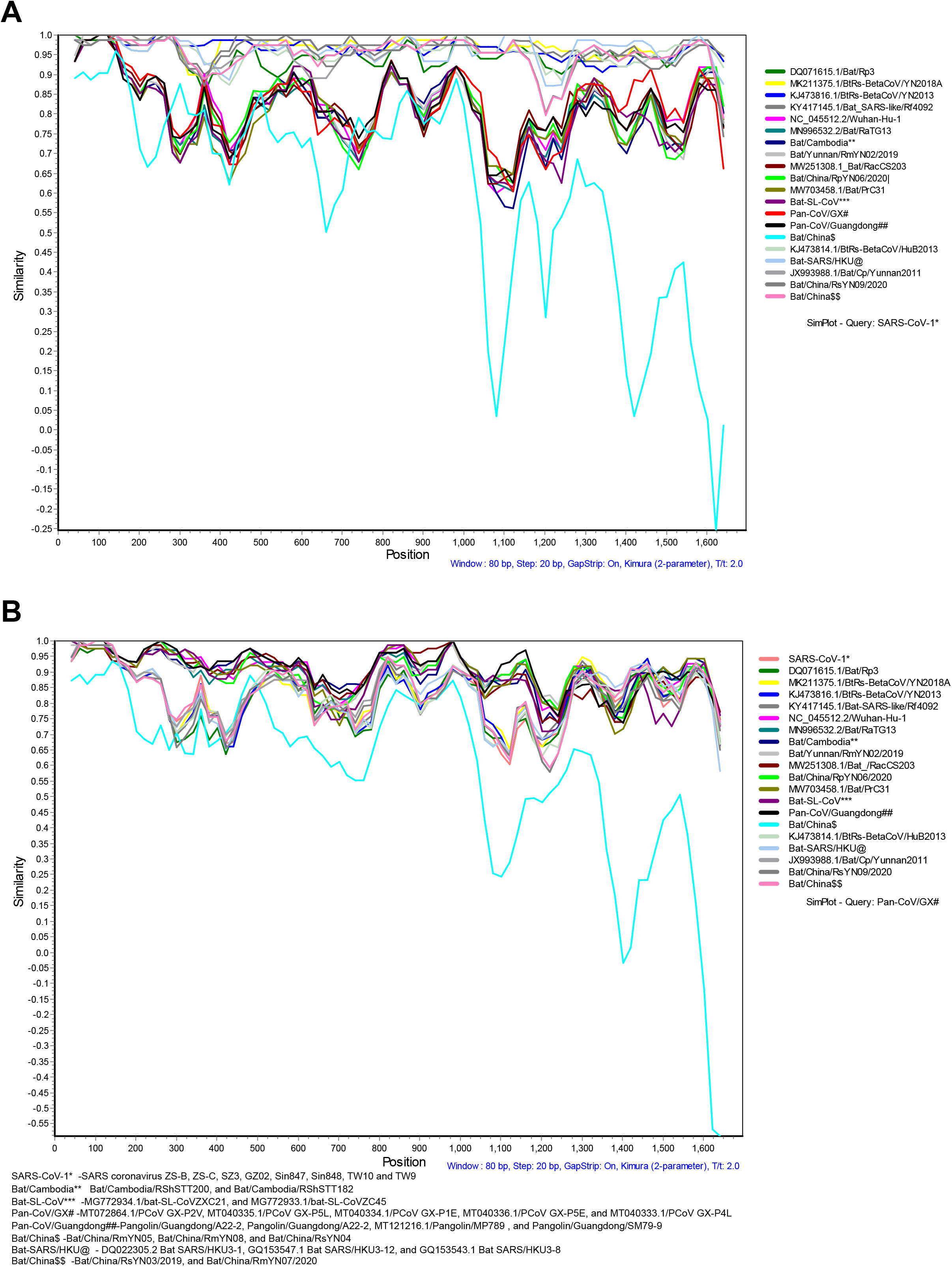
Similarity plot analysis for the E, M, protein 6, protein 7a, and protein 7b genes. (**A**) Similarity plot analysis for the Pan-CoVs, SARS-CoV-2, and related viruses using the nucleotide sequences of E, M, protein 6, protein 7a, and protein 7b.The SARS-CoV-1 is set as a query sequence. (**B**) Similarity plot analysis for the Pan-CoVs, SARS-CoV-2, and related viruses using the nucleotide sequences of E, M, protein 6, protein 7a, and protein 7b. The Pan-CoV/GX lineages are set as a query sequence.

**Supplementary Figure 6.**
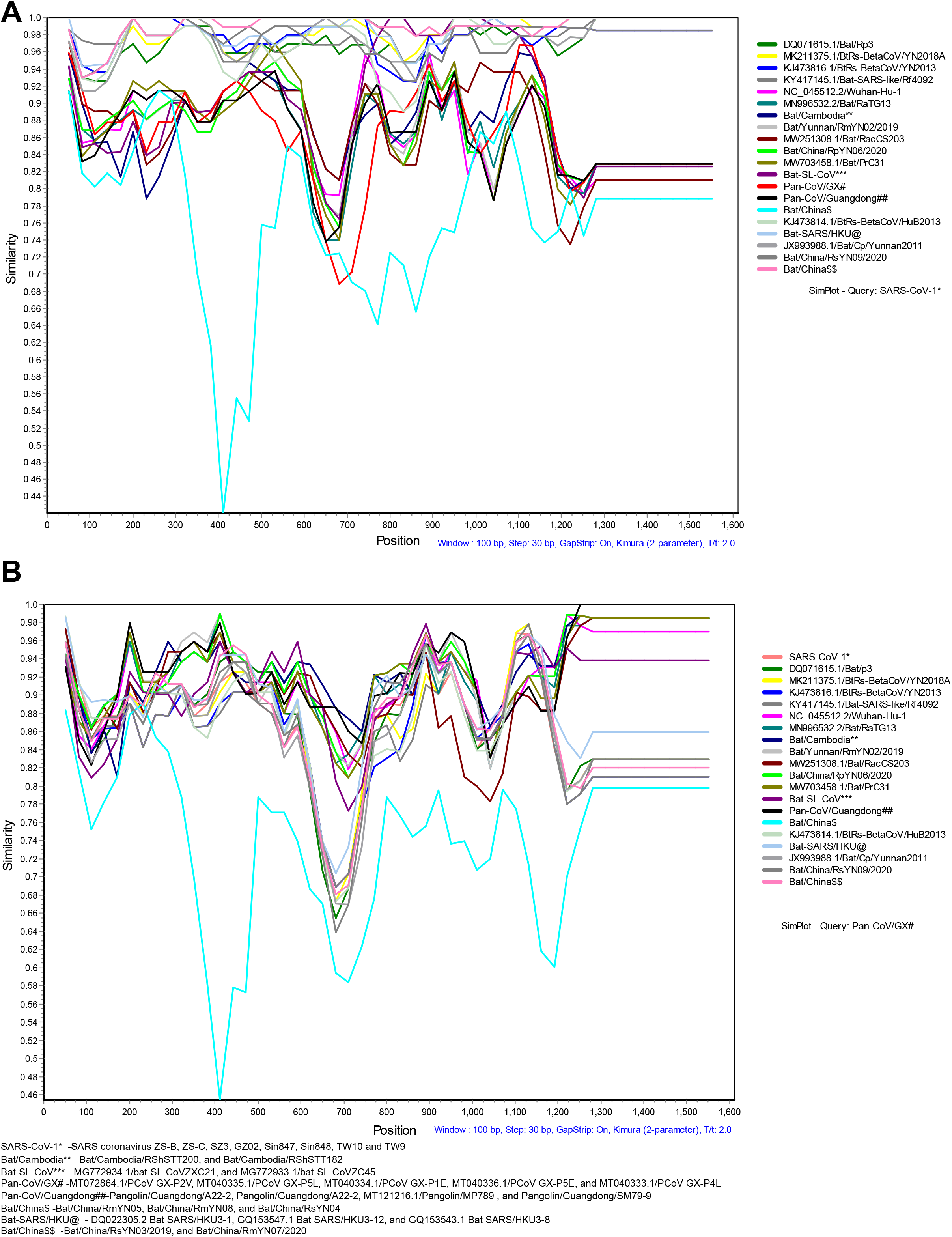
Similarity plot analysis for the complete nucleocapsid (N), ORF10, and 3’UTR. (**A**) Similarity plot for the Pan-CoVs, SARS-CoV-2, and related viruses using the nucleotide sequences of the nucleocapsid (N), ORF10, and 3’UTR. The SARS-CoV-1 is set as a query sequence. (**B**) Similarity plot for the Pan-CoVs, SARS-CoV-2, and related viruses using the nucleotide sequences of the nucleocapsid (N), ORF10, and 3’UTR. The Pan-CoV/GX lineages are set as a query sequence.

**Supplementary Figure 7.**
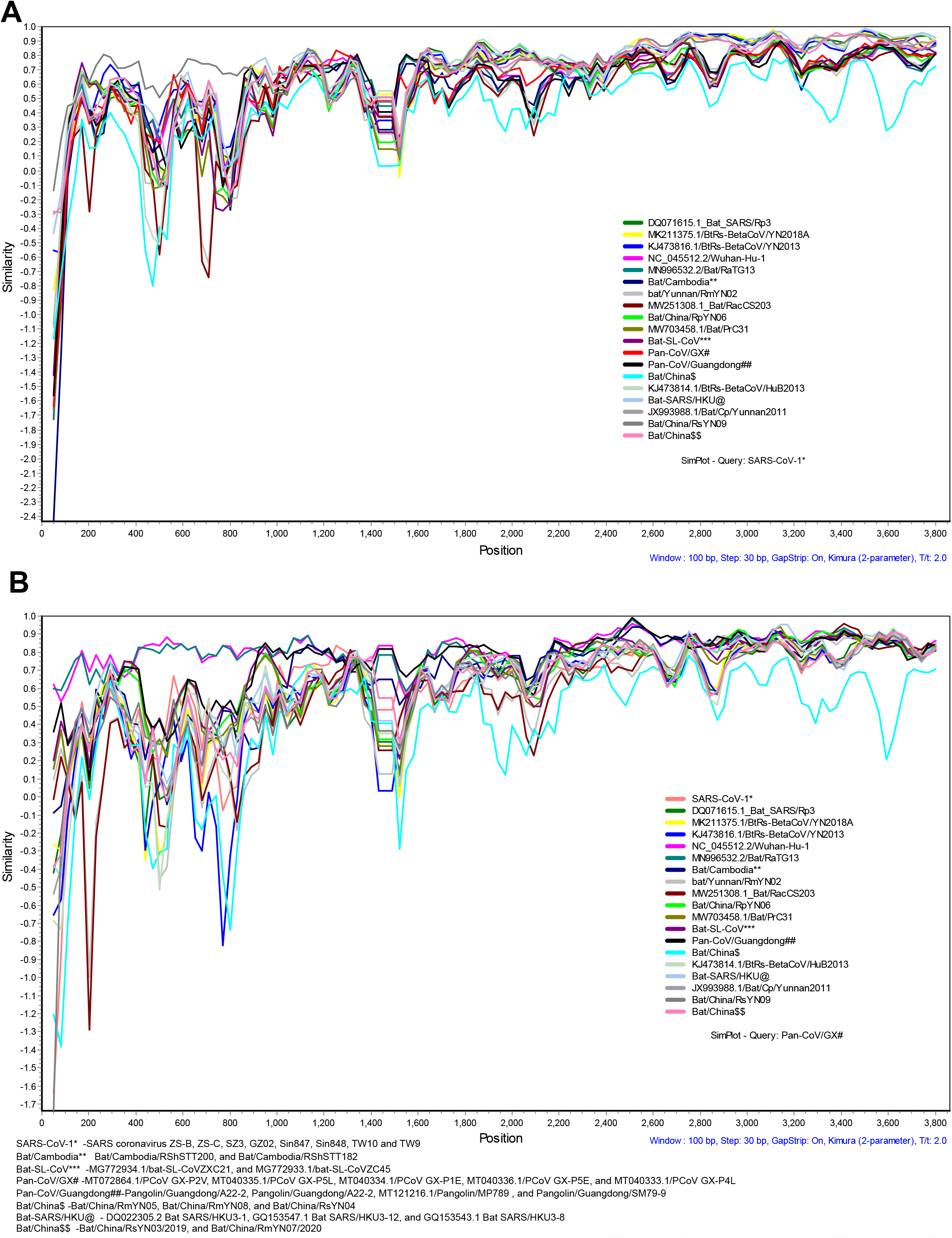
Similarity plot analysis for the complete spike gene. (**A**) Similarity plot for the Pan-CoVs, SARS-CoV-2, and related viruses using the nucleotide sequences of complete spike gene. The SARS-CoV-1 is set as a query sequence. (**B**) Similarity plot for the Pan-CoVs, SARS-CoV-2, and related viruses using the nucleotide sequences of complete spike gene. The Pan-CoV/GX lineages are set as a query sequence.

